# Distinctive alteration of presynaptic proteins in the outer molecular layer of the dentate gyrus in Alzheimer’s disease

**DOI:** 10.1101/2020.10.06.327833

**Authors:** Hazal Haytural, Tomás Jordá-Siquer, Bengt Winblad, Christophe Mulle, Lars O. Tjernberg, Ann-Charlotte Granholm, Susanne Frykman, Gaël Barthet

**Affiliations:** Division of Neurogeriatrics, Center for Alzheimer Research, Department of Neurobiology, Care Sciences and Society, Karolinska Institutet, Solna, Sweden; Univ. Bordeaux, CNRS, Interdisciplinary Institute for Neuroscience, IINS, UMR 5297, F-33000 Bordeaux, France; Karolinska University Hospital, Theme Aging, Huddinge, Sweden; Knoebel Institute for Healthy Aging, University of Denver, Denver, Colorado

**Keywords:** Alzheimer’s Disease, Postmortem human brain, Hippocampus, Outer molecular layer of Dentate Gyrus, Perforant path, Presynaptic impairment, Immunofluorescence

## Abstract

Synaptic degeneration has been reported as one of the best pathological correlate of cognitive deficit in Alzheimer’s Disease (AD). However, the location of these synaptic alterations within hippocampal sub-regions, the vulnerability of the presynaptic versus postsynaptic compartments, and the biological mechanisms for these impairments remain unknown. Here, we performed immunofluorescence labeling of different synaptic proteins in fixed and paraffin embedded human hippocampal sections and report reduced levels of several presynaptic proteins of the neurotransmitter release machinery (complexin-1, syntaxin-1A, synaptotagmin-1 and synaptogyrin-1) in AD cases. The deficit was restricted to the outer molecular layer (OML) of the dentate gyrus whereas other hippocampal sub-fields were preserved. Interestingly, standard markers of postsynaptic densities (SHANK2) and dendrites (MAP2) were unaltered, as well as the relative number of granule cells in the dentate gyrus, indicating that the deficit is preferentially presynaptic. Notably, staining for the axonal components, myelin basic protein, SMI-312 and Tau, was unaffected, suggesting that the local presynaptic impairment does not result from axonal loss or alterations of structural proteins of axons. There was no correlation between the reduction in presynaptic proteins in OML and the extent of the amyloid load or of the dystrophic neurites expressing phosphorylated forms of Tau. Altogether, this study highlights the distinctive vulnerability of the OML of dentate gyrus and supports the notion of presynaptic failure in AD.

## Introduction

Synaptic dysfunction and degeneration are early features in Alzheimer’s Disease (AD) pathogenesis and correlate well with measures of cognitive decline [11, 29, 53]. The hippocampus, a brain region which is essential for episodic memory formation is severely impaired in AD. The hippocampus consists of different anatomical regions including the dentate gyrus, the cornu ammonis (CA) regions, and the subiculum [33] (Fig. 1a). The main input to the hippocampus comes from the entorhinal cortex (EC) through the perforant path [9] (Fig. 1a, c). The neurons from the layer II of the EC project to the outer two-thirds of the molecular layer of the dentate gyrus (hereafter referred as to outer molecular layer, OML; Fig. 1c) as well as to CA3, whereas the neurons from layer III project to CA1 and subiculum regions (Table 1). The hippocampus also receives major modulatory input from the serotonergic, noradrenergic, and dopaminergic transmitter systems as well as from the medial septal nucleus (cholinergic neurons) via the fimbria/fornix pathway [47]. The perforant path has been proposed to be vulnerable in AD pathogenesis due to (i) synaptic loss observed in the OML of AD and mild cognitive impairment cases [43, 44]; (ii) the presence of AD-hallmarks, such as amyloid plaques and neurofibrillary tangles, both in the EC [2, 6, 54] and in the OML [22, 54]; and (iii) substantial loss of EC neurons in AD cases, particularly of layer II [16, 24, 36], as well as loss of cholinergic input to the hippocampus [17].

**Table 1.** Summary of afferents and efferents of the hippocampus.

| Afferents |  | Efferents |  |
| --- | --- | --- | --- |
| **OML** | <sup>1</sup> Perforant path fibers from entorhinal cortex layer II | **Granule cell layer** | <sup>1</sup> CA3 via mossy fibers |
| **IML** | <sup>1</sup> Associational/Commissural fibers from hilus<br><sup>2</sup> CA3 collaterals<br><sup>3</sup> Septal fibers<br><sup>4</sup> Fibers from thalamus |  |  |
| **CA3-LUC** | <sup>1</sup> Mossy fibers from hilus<br><sup>2</sup> Fibers from locus coeruleus | **CA3** | <sup>1</sup> CA1 via Schaffer collaterals |
| **CA3-RAD** | <sup>1</sup> CA3 associational fibers<br><sup>2</sup> Septal fibers |  |  |
| **CA3-LM** | <sup>1</sup> Perforant path fibers from entorhinal cortex layer II<br><sup>2</sup> Septal fibers<br><sup>3</sup> Fibers from locus coeruleus |  |  |
| **CA1-RAD** | <sup>1</sup> Schaffer collaterals from CA3<br><sup>2</sup> Recurrent collaterals from CA1<br><sup>3</sup> Fibers from amygdaloid complex | **CA1** | <sup>1</sup> Entorhinal cortex layer V via Subiculum |
| **CA1-LM** | <sup>1</sup> Perforant path fibers from entorhinal cortex layer III<br><sup>2</sup> Fibers from thalamus |  |  |
|  | <sup>3</sup> Fibers from amygdaloid complex |  |  |

**Figure 1.**
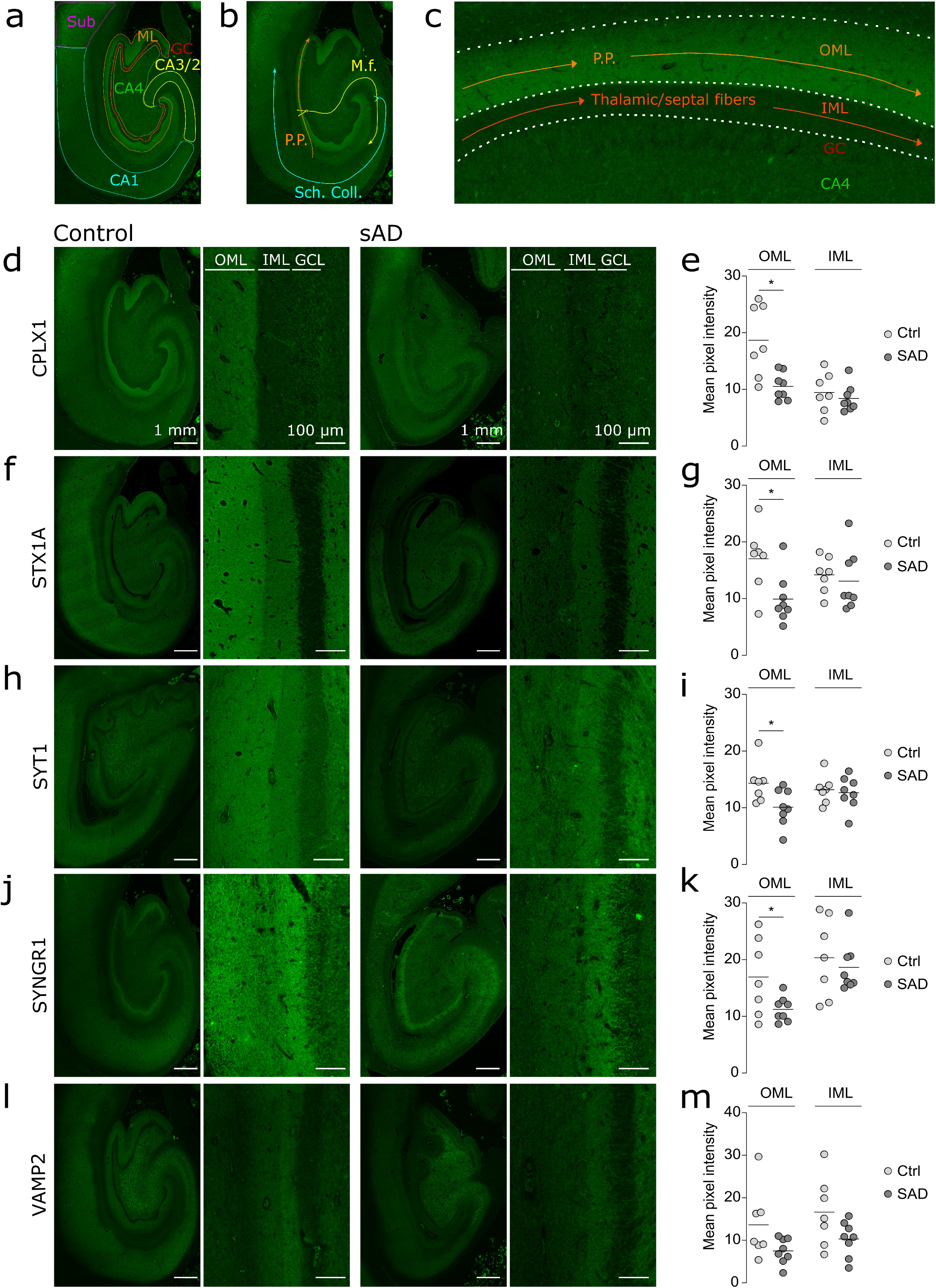
OML-specific reduction in presynaptic proteins. (a) Schematic diagram showing the main hippocampal sub-regions in a section labelled for CPLX1 in a healthy hippocampus. (b) Schematic diagram showing the hippocampal trisynaptic circuit. (c) Schematic diagram showing the main afferents projections in OML and IML. (d, f, h, i, j) Representative images of the whole hippocampus and zoom-in pictures of the molecular region of the DG in controls (left) and AD (right) brain sections labelled respectively for CPLX1, STX1A, SYT1, SYNGR1, VAMP2. The regions of interests are indicated by outer (OML) and inner (IML) molecular layers, which are located right next to the granule cell layer (GCL). (e, g, i, k, m) Scatter plots of the mean pixel intensity of the fluorescent labelling in OML and IML respectively for CPLX1, STX1A, SYT1, SYNGR1, VAMP2. The fluorescence pixel intensities were significantly decreased in AD cases by 44% for CPLX1 (c, p-value = 0.0051), 42 % for STX1A (d, p-value = 0.016), 29 % for SYT1 (e, p-value = 0.031) and 34% for SYNGR1 (f, p-value = 0.042). In contrast, the levels of these proteins were not altered in the IML of AD cases. (m) The levels of VAMP2 showed a non-significant reduction by 45% (p-value = 0.068) and 38% (p-value = 0.075) both in the OML and the IML, respectively. Note that in a healthy brain, CPLX1, STX1A and SYT1 displayed a stronger IR in the OML with respect to IML (d, f, h), while SYNGR1- and VAMP2-IR was more prominent in the IML than the OML (j, l). Scale bar is 1 mm (low magnification) and 100 μm (high magnification).

In order to further delineate the synaptic pathology of OML in AD brain, we recently performed a proteomic study of microdissected OML from five AD (Braak IV and amyloid C) and five control cases [18]. We found that a large number of presynaptic proteins were significantly decreased in the OML of AD brains, whereas postsynaptic proteins were relatively spared. In addition, extensive pathway analysis indicated that synaptic vesicle cycling and exocytosis as prominently affected pathways in the OML of AD brains. The current work is an extension of this earlier publication, where we investigate the specificity and the mechanisms leading to the alteration of presynaptic proteins by performing immunofluorescent stainings in hippocampal sections from human AD and control cases. We selected five synaptic proteins: complexin-1 (CPLX1), synaptotagmin-1 (SYT1), syntaxin-1A (STX1A), synaptogyrin-1 (SYNGR1) and vesicle-associated membrane protein 2 (VAMP2), based on the extent of their reduction determined in our proteomic assay [17], their presynaptic function and the availability of validated antibodies.

STX1A, SYT1 and VAMP2 are all part of the core synaptic vesicle-membrane-fusion machinery and thus play important roles in neurotransmission [49]. STX1A together with the synaptic vesicle protein VAMP2 and the plasma membrane protein SNAP-25 form the soluble NSF-attachment protein receptor (SNARE) complex [48, 50], which is the key component of membrane fusion machinery in exocytosis [49]. SYT1 is a synaptic vesicle protein that acts as a Ca^2+^ sensor [15]; increased levels of Ca^2+^ in the presynaptic terminal triggers an interaction between SYT1 and the plasma membrane protein STX1A that is important for synaptic vesicle exocytosis [45]. Moreover, complexins, such as CPLX1, bind to SNARE complexes [31], participate in Ca^2+^-triggered exocytosis [38], and therefore are essential for neurotransmitter release [28]. Among these proteins, the synaptic vesicle protein SYNGR1 is the least studied. Although its exact function still remains unknown, earlier studies suggest that SYNGR1 is most likely not essential for exocytosis, but it may play a regulatory role in synaptic plasticity, and therefore, neurotransmitter release [23].

Although decreased mRNA and/or protein levels of these five presynaptic markers have been previously detected in AD brain [4, 7, 32, 51], these studies have so far been performed on homogenates prepared from cortical or hippocampal bulk tissue, thus not allowing any detailed examination of different hippocampal sub-regions. Here, we investigated the distribution of presynaptic proteins in 10 hippocampal sub-regions of AD and control cases using immunofluorescence. We observed a reduction of presynaptic protein staining densities in the OML consistent with the reduction determined in our proteomic approach. Interestingly, this reduction appeared to be specific to the OML since all others hippocampal sub-regions examined, displayed a similar expression level between controls and AD cases. In order to depict what underlying causes might be responsible for the observed changes in the presynaptic proteins in AD OML, we further investigated both post- and presynaptic compartments by assessing: (1) staining density of postsynaptic protein and a dendritic marker, (2) the relative number of dentate granule cells, and finally (3) the staining densities of axonal markers. Finally, we investigated a possible mechanisms involving the major AD hallmarks, amyloid plaques and phosphorylated-Tau.

## Results

We investigated the specificity and the mechanisms leading to the alteration of presynaptic proteins in AD by comparing hippocampus sections from AD (n=8) and control (n=7) patients from the Netherlands Brain Bank. There were no statistically significant differences in gender, age, postmortem interval (PMI), ApoE status, or brain pH between the two groups, while Braak and amyloid stages showed statistically significant differences, as expected (Table 2).

**Table 2.** Clinical and neuropathological details of cases.

| Case | Diagnosis | Gender | Age | PMI (h) | Brain pH | APOE status | Braak tau stage | Amyloid stage |
| --- | --- | --- | --- | --- | --- | --- | --- | --- |
| 1 | C | F | 73 | 7.5 | n/a | 44 | 1 | B |
| 2 | C | F | 84 | 6 | 7.65 | 33 | 2 | B |
| 3 | C | F | 76 | 7 | 6.87 | 33 | 2 | O |
| 4 | C | F | 81 | 5.5 | 6.77 | 33 | 3 | A |
| 5 | C | F | 84 | 5.5 | 6.68 | 33 | 2 | A |
| 6 | C | F | 83 | 6 | 6.6 | 44 | 2 | B |
| 7 | C | F | 82 | 5 | 6.34 | 33 | 2 | B |
| 8 | AD | F | 85 | 6 | 6.35 | 44 | 5 | C |
| 9 | AD | F | 90 | 4.5 | 6.39 | 33 | 6 | C |
| 10 | AD | F | 70 | 5 | 6.73 | 33 | 5 | C |
| 11 | AD | F | 91 | 6.5 | 6.01 | 44 | 5 | C |
| 12 | AD | F | 70 | 6 | 6 | 33 | 6 | C |
| 13 | AD | F | 80 | 4 | 6.26 | 44 | 6 | C |
| 14 | AD | F | 70 | 4 | 6.42 | 44 | 6 | C |
| 15 | AD | F | 86 | 5 | 6.2 | 33 | 6 | C |
| p-value: |  | NS | NS | NS | NS | NS | 0.0002 | 0.0002 |

The clinical and pathological characteristics were compared between AD and control groups using either Student's t-test (age, PMI and brain pH) or Mann Whitney test (gender, ApoE status, Braak and CERAD scores), and p-value < 0.05 considered statistically significant.

We used an immunofluorescence approach combining an antigen retrieval and autofluorescence quenching methods (see absence of auto-fluorescence in controls; Supplementary Fig. 1a) to label synaptic proteins in in fixed, paraffin embedded sections of human brain tissue. The mean fluorescence pixel intensity of synaptic proteins in different anatomical regions was measured to achieve semi-quantitative analysis of protein abundance.

### Marked reduction of presynaptic proteins levels in the OML in AD

The fluorescence pixel intensities of the five presynaptic proteins CPLX1, STX1A, SYT1, SYNGR1 and VAMP2 were measured in the OML and inner molecular layer (IML) of the dentate gyrus. Interestingly, we found that these proteins displayed different spatial distribution between IML and OML. In a control brain, staining with antibodies directed against CPLX1, STX1A and SYT1 gave rise to a stronger immuno-reactivity (IR) in OML than in IML (Fig. 1d-i), while SYNGR1 and VAMP2 appeared to be more abundantly expressed in IML than in OML (Fig. 1j-m). The region-specific distribution of these presynaptic proteins created a visible line in the molecular layer, thus allowing us to easily separate IML and OML by gross visualization (Figure 1).

Semi-quantitative densitometric analysis showed that the staining densities of CPLX1 (44 % decrease, p-value = 0.0051, Fig. 1d, e), STX1A (42 % decrease, p-value = 0.016, Fig. 1f, g), SYT1 (29 % decrease, p-value = 0.031, Fig. 1h, i), and SYNGR1 (34 % decrease, p-value = 0.042, Fig. 1j, k) were significantly reduced in OML in AD compared to control cases, whereas there was a non-significant decreased tendency in the same direction for VAMP2 staining (45 % decrease, p-value = 0.068, Fig. 1l, m).

Interestingly, we observed no profound changes between the two groups in the levels of these presynaptic proteins in the IML which receives afferents from the medial septum [4], the thalamus [6] and the hilus of the dentate gyrus [2] (Fig. 1d-m). When the mean pixel intensity of a synaptic protein in OML was normalized to the intensity of the same protein in IML, the ratio was significantly decreased for CPLX1, STX1A, SYT1 and SYNGR1 but not for VAMP2 (Supplementary Fig. 1) giving further support to the notion of a specific impairment of OML compared to IML.

### Presynaptic protein staining reduction in AD hippocampus specific to the OML

The marked difference in the reduction of presynaptic proteins in OML compared to IML indicates that the synaptic alterations depend on the afferent inputs within a brain sub-region. This led us to investigate whether other hippocampal sub-regions could be affected – which might be innervated by other modalities. The OML receiving abundant afferents from the EC, we hypothesized that other hippocampal sub-regions receiving EC projections (Table 1) could be affected. Thus, in addition to the OML and IML, we analyzed the fluorescence pixel intensities of CPLX1, STX1A, SYT1, SYNGR1 and VAMP2 in 8 different sub-regions of the hippocampus visible on the same sections, categorized into two groups: (i) molecular layers, enriched in synapses, consisting of CA3-LUC, CA3-RAD, CA3-LM, CA1-RAD and CA1-LM, and (ii) neuronal layers, consisting of CA4, CA3 and CA1 neuronal layers. Despite the observed reduction in CPLX1, SYT1, SYNGR1 levels in AD OML, no significant reductions were detected in any other investigated molecular layers in AD compared to control cases (Fig. 2a-j) or between neuronal layers (see pictures in Fig. 1 and quantifications in Supplementary Fig. 2). Interestingly, the cell-surface SNARE protein STX1A exhibited a markedly different staining profile in AD than the vesicular or cytosolic presynaptic proteins described above. Indeed, a marked increase in STX1A staining was observed in the CA4 neuronal layer of AD cases compared to controls (36 % increase, p-value = 0.048, Supplementary Fig. 2b). Moreover, non-significant trends towards increased levels of STX1A were detected also in the CA3 neuronal layer (52 % increase, p-value = 0.0506), CA3-LUC (55 % increase, p-value = 0.08), CA3-RAD (34 % increase), CA1 neuronal layer (18 % increase), and CA1-RAD (31 % increase, p-value = 0.09) (molecular layers in Fig. 2b; neuronal layers in Supplementary Fig. 2b).

**Figure 2.**
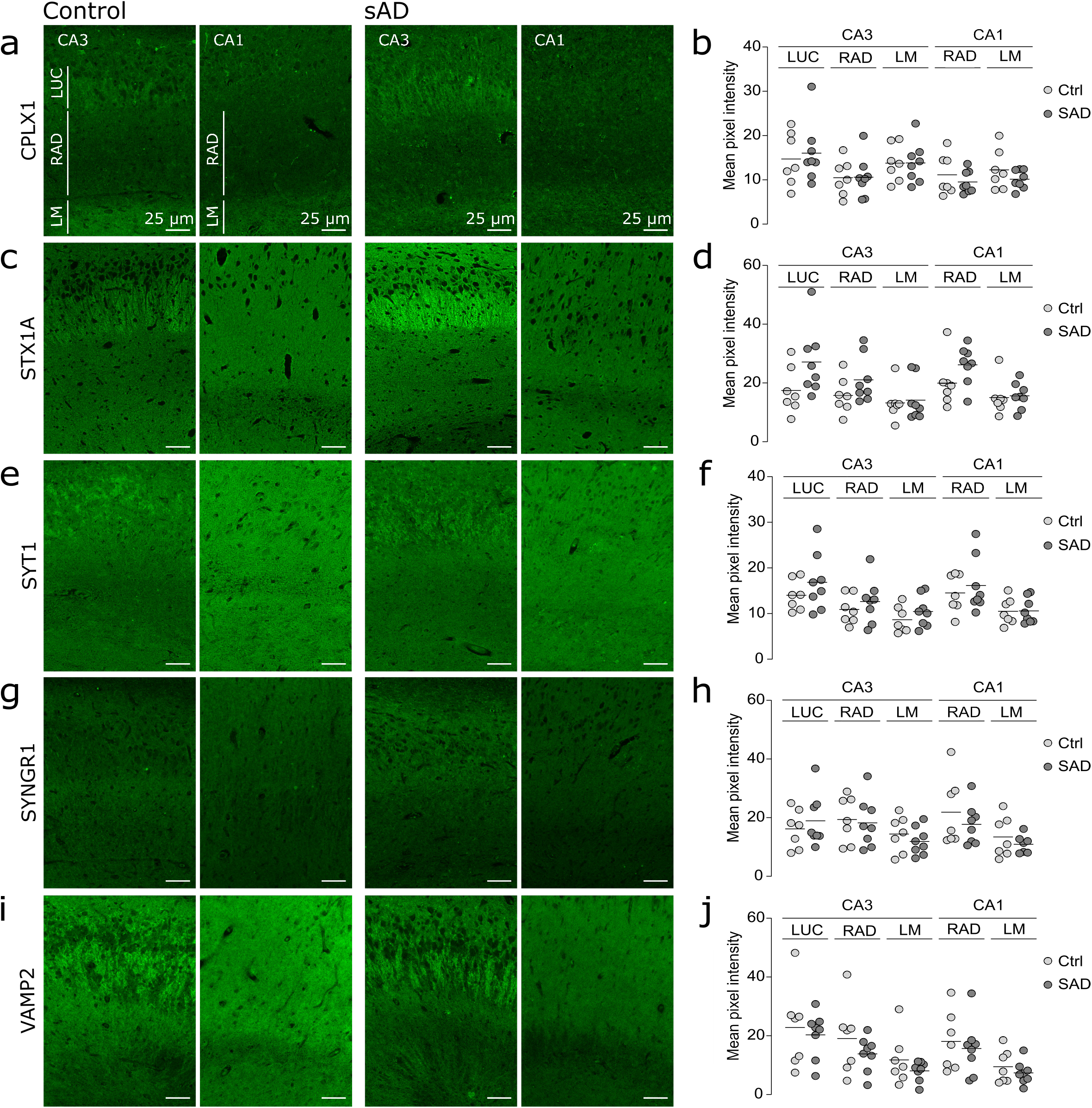
Preserved level of presynaptic proteins in various molecular layers of the hippocampus. (a, c, e, g, i) Representative images of the CA3 and CA1 regions in controls (left) and AD (right) brain sections labelled respectively for CPLX1, STX1A, SYT1, SYNGR1, VAMP2. The region of interests stratume lucidum (LUC), radiatum (RAD) and lacunosum moleculare (LM) are indicated. (b, d, f, h, j) Scatter plots of the mean pixel intensity of the fluorescent labelling in OML and IML respectively for CPLX1, STX1A, SYT1, SYNGR1, VAMP2. The mean pixel intensities were not altered between AD and control groups in the molecular layers of the CA3 and CA1. Scale bar is 25 μm.

### Preserved postsynaptic compartment in OML of AD cases

The observation of reduced levels of presynaptic proteins in OML raised the question of whether the postsynaptic proteins were also reduced, as expected from the hypothesis of an overall synaptic loss. We assessed the levels of the standard markers of postsynaptic density and dendrites, respectively SHANK2 and MAP2. Despite the presence of severe Braak and amyloid stages among the AD cases, the levels of SHANK2 and MAP2 were not reduced in AD compared to controls (Fig. 3a-d). Furthermore, we investigated the density of dentate granule cells, the dendrites of which are located at the OML forming synapses with the perforant path termini and noted that there was no obvious reduction in granule cells density in AD compared to control cases (Fig. 3e, f). Together, these results advocate for a distinctive impairment of presynaptic proteins in OML synaptic inputs rather than an overall synaptic loss.

**Figure 3.**
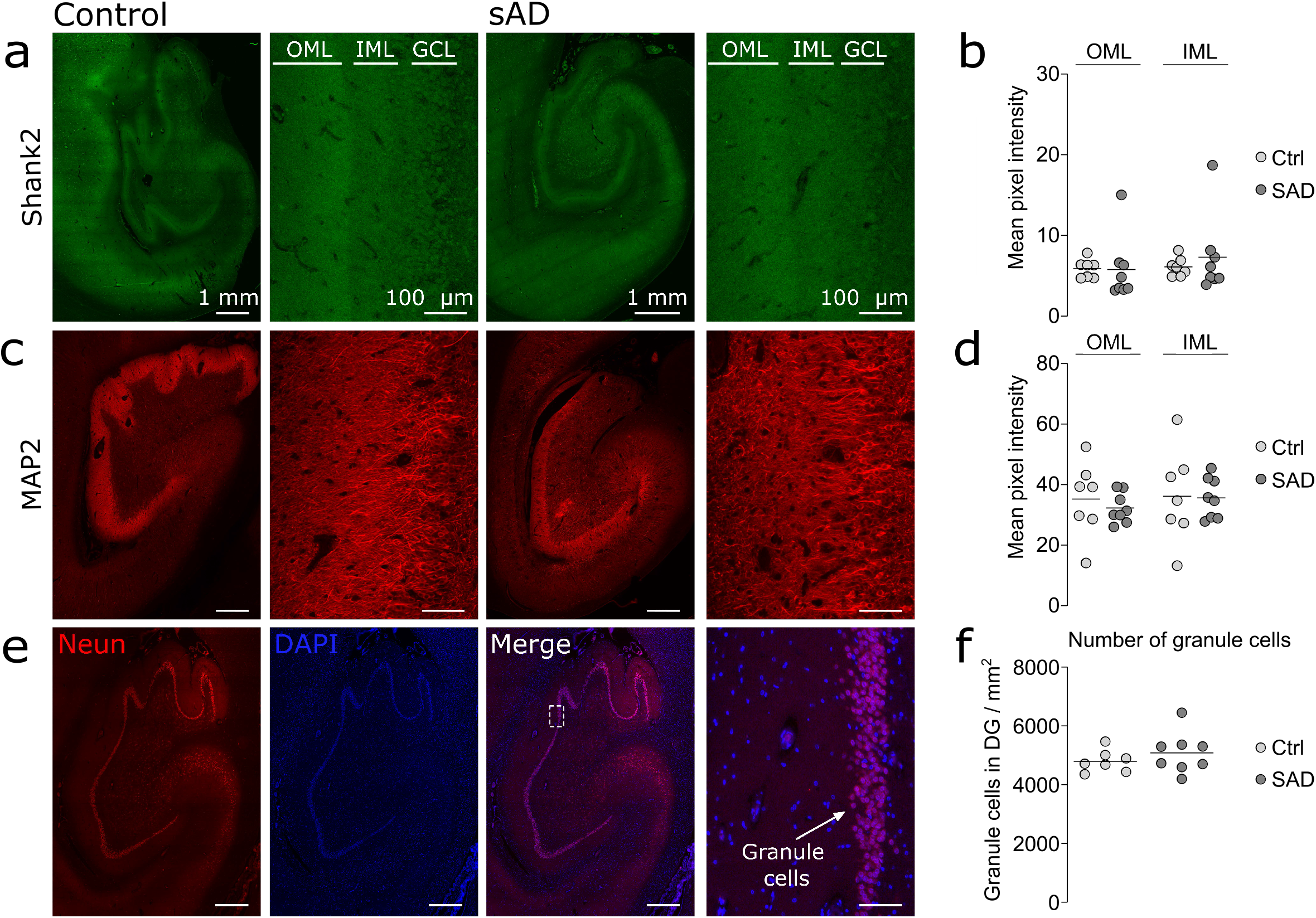
Preserved postsynaptic compartment in AD OML. (a, c, e) Representative images of the whole hippocampus and zoom-in pictures of the molecular region of the DG in controls (left) and AD (right) brain sections labelled respectively for Shank2, MAP2 and colabelled for NeuN and DAPI. The regions of interests are indicated by inner (IML) and outer (OML) molecular layers, which are located right next to the granule cell layer (GCL). Scale bars are 1 mm (low magnification) and 100 μm (high magnification). (b, d) Scatter plots of the mean pixel intensity of the fluorescent labellings in OML and IML respectively for Shank2 and MAP2. The levels of SHANK2 and MAP2 were not altered. (f) Scatter plots of the mean pixel intensity of the neuronal density (number of dentate granule cells per mm^2^) did not show a difference between AD and control groups.

### Evidence for maintained axonal projections in AD OML

We next investigated potential mechanisms involved in alterations of presynaptic proteins in OML in AD. One potential mechanisms would be if a reduction of axonal projections to the OML would proportionally lead to a reduced amount of presynaptic proteins in this region. Consecutive hippocampal sections from AD and control cases were stained with three different axonal markers: myelin basic protein (MBP; for myelinated axons), SMI-312 (for phosphorylated neurofilaments M and H) and the commonly used axonal marker total Tau. Semi-quantitative densitometric assessment showed no difference in the levels of above-mentioned axonal markers between the two groups (Fig. 4a-f) suggesting that biological mechanisms other that a reduction in afferent fibers density are at play in the alterations in presynaptic proteins in OML.

**Figure 4.**
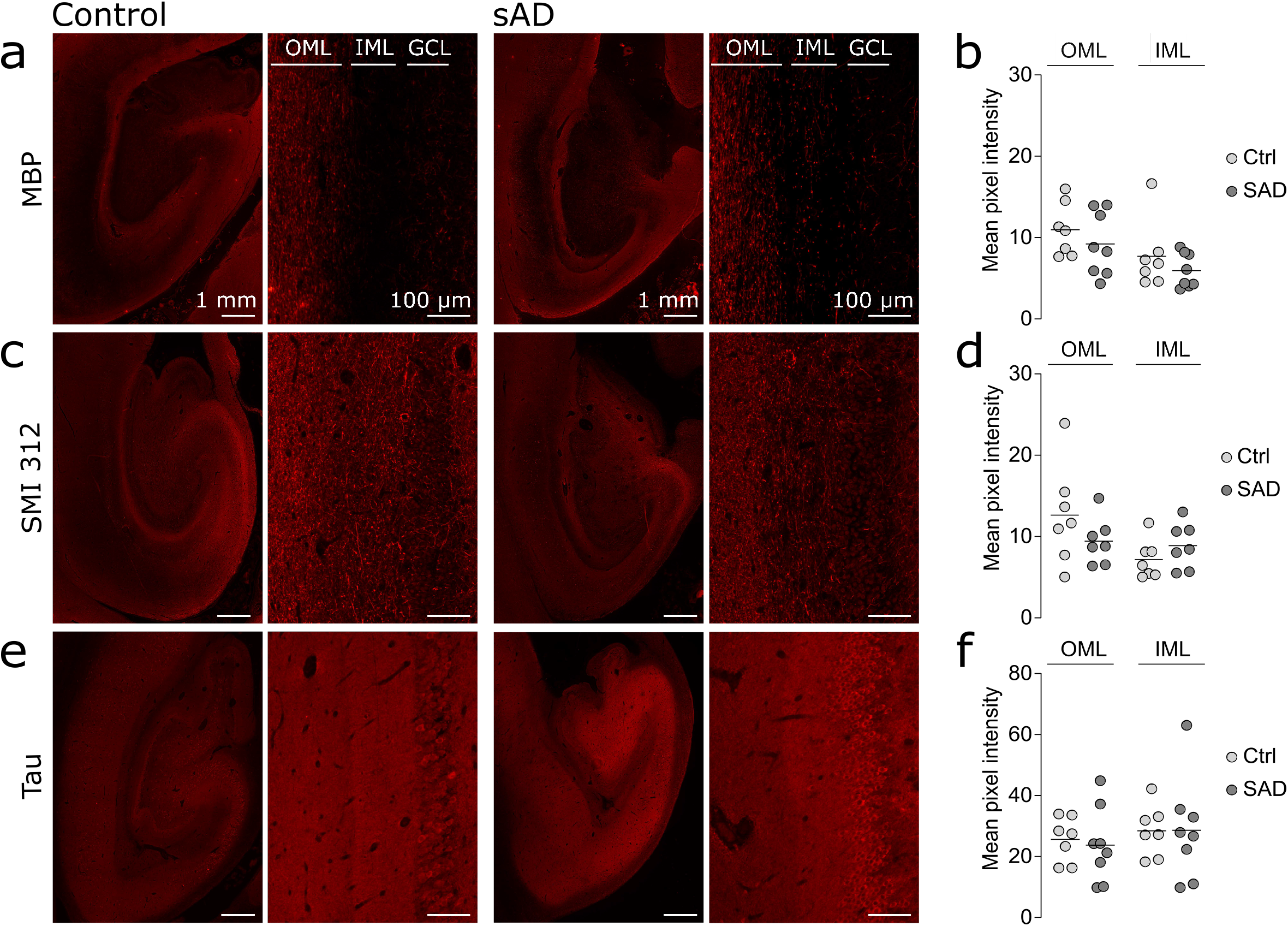
Evidence for maintained axonal projections in AD OML. (a, c, e) Representative images of the whole hippocampus and zoom-in pictures of the molecular region of the DG in controls (left) and AD (right) brain sections labelled respectively for MBP, SMI 312 and Tau. The regions of interests are indicated by outer (OML) and inner (IML) molecular layers, which are located right next to the granule cell layer (GCL). Scale bars are 1 mm (low magnification) and 100 μm (high magnification). (b, d, f) Scatter plots of the mean pixel intensity of the fluorescent labelling in OML and IML respectively for MBP, SMI 312 and Tau. The levels of MBP, SMI 312 and Tau were not altered.

### Presynaptic failure in OML not related to AD pathology

To further explore the mechanisms behind the specific impairment in presynaptic proteins in OML, we investigated the presence of AD-related pathological hallmarks in the different hippocampal sub-regions of interest. We immuno-labelled amyloid plaques with an antibody against Aβ peptides (clone 6E10) and pathological forms of Tau with an antibody against its phosphorylated isoforms (AT8). Dense core amyloid plaques were distinctly observable in AD cases whereas they were absent from the controls (Fig. 5a). The extent of the amyloid burden defined as the total surface covered by Aβ peptides normalized to the surface of the sub-regions analyzed was not significantly different between OML and the other hippocampal regions (Fig. 5b). Besides, within OML there was no correlation between amyloid load and the extent of presynaptic proteins reduction among the different AD patients (Fig. 5c). Together, these data indicate that the distinctive reduction in presynaptic proteins in OML cannot be explained by a larger amyloid burden in this sub-region.

**Figure 5.**
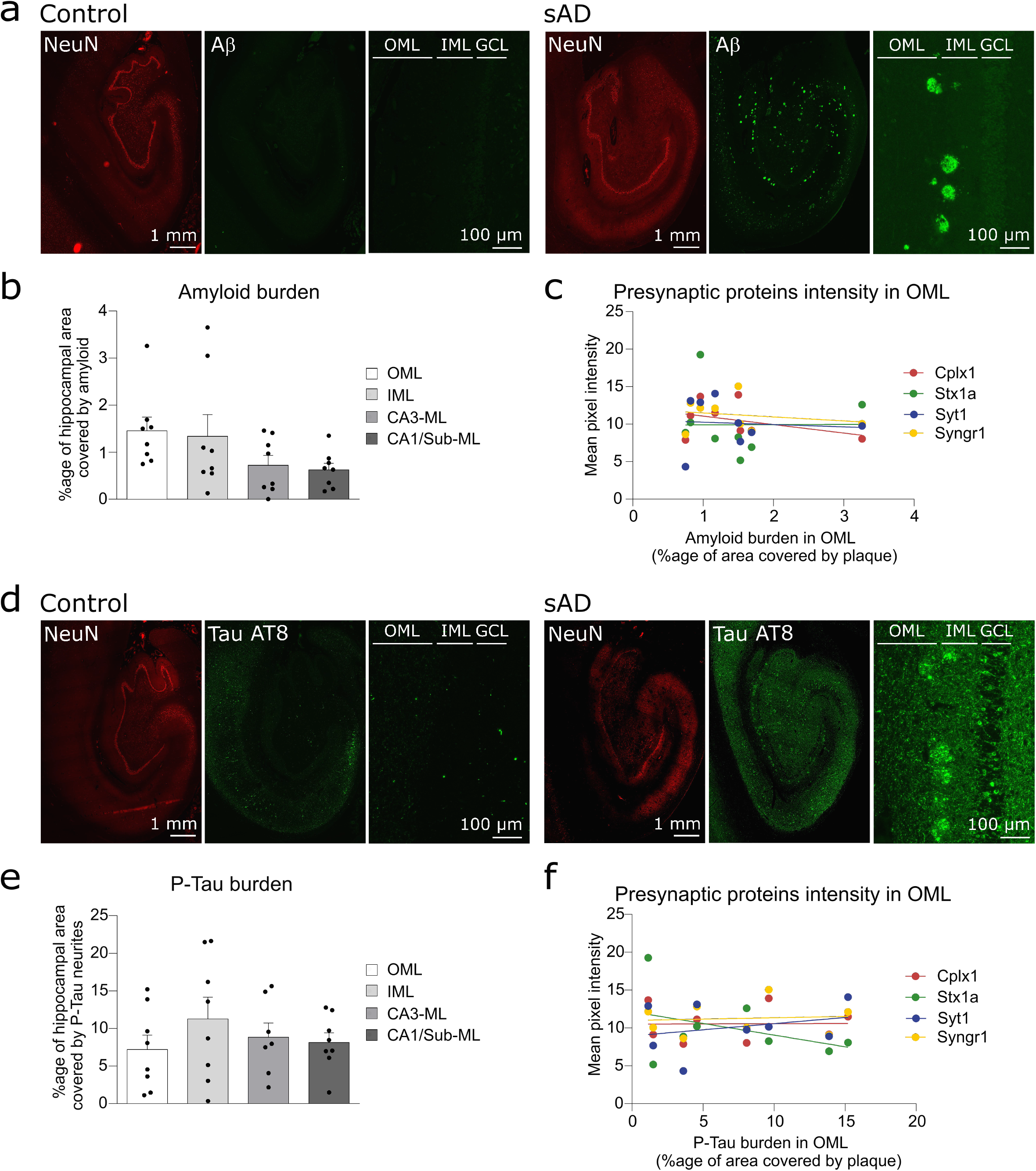
AD-related hallmarks relationship to presynaptic proteins decrease. (a) Representative images of the whole hippocampus and zoom-in pictures of the molecular region of the DG in controls (left) and AD (right) brain sections labelled respectively for NeuN and Aβ1-16. The regions of interests are indicated by outer (OML) and inner (IML) molecular layers located next to the granule cell layer (GCL). Scale bars are 1 mm (low magnification) and 100 μm (high magnification). (b) Scatter plots with bars of the mean percentage of area covered by amyloid staining in different hippocampal molecular layers (OML, IML, CA3-ML, CA1-ML). The amyloid burden is not significantly different between the regions. (c) Graph of the mean pixel intensity of the presynaptic protein CPLX1, STX1A, SYT1 and SYNGR1 in OML in function of the amyloid burden. Each dot represents the value of one AD case. There was no correlation between the amyloid burden and the extent of the presynaptic proteins reduction. (d) Representative images of the whole hippocampus and zoom-in pictures of the molecular region of the DG in controls (left) and AD (right) brain sections labelled respectively for NeuN and phosphorylated-Tau (AT8). The regions of interests are indicated by outer (OML) and inner (IML) molecular layers located next to the granule cell layer (GCL). Scale bars are 1 mm (low magnification) and 100 μm (high magnification). (e) Scatter plots with bars of the mean percentage of area covered by neurites positive for phosphorylated-Tau (AT8) in different hippocampal subregions (OML, IML, CA3, CA1). The dystrophic neurites burden is not significantly different between the regions. (f) Graph of the mean pixel intensity of the presynaptic protein CPLX1, STX1A, SYT1 and SYNGR1 in OML in function of the burden in AT8-positive dystrophic neurites. Each dot represents the value in one AD case. There was no correlation between the burden in AT8-positive dystrophic neurites and the extent of the presynaptic proteins reduction.

The antibody against phosporylated Tau (AT8) labelled numerous soma in AD cases, in the different CA regions and in the dentate gyrus (Fig. 5d). Strikingly, the abundance of Tau (AT8)-positive neurites was massively higher in AD compared to controls (Fig. 5d). We hypothesized that a higher density of dystrophic neurites, as revealed by phosphorylated Tau (AT8) immunogenicity, could be related to the presynaptic impairment. However, the extent of the burden in dystrophic neurites, i.e. the total surface covered by dystrophic neurites positive for phosphorylated Tau (AT8), normalized to the surface of the sub-region analyzed was not significantly different between OML and the other hippocampal molecular layers (Fig. 5e). Finally, within OML there was no correlation between the burden in dystrophic neurites and the extent of presynaptic proteins reduction among the different AD patients (Fig. 5c).

## Discussion

Synaptic dysfunction, as revealed by reduction in synaptic proteins [56] as well as loss of synapses [41], have been observed in early stages of AD. To have a better insight into the molecular mechanisms involved, we previously performed a proteomic analysis of the OML, one of the terminal zones of the perforant path, the main afferent pathway innervating the hippocampus. This approach revealed an impairment of OML proteome in AD characterized by a reduced level of presynaptic proteins whereas the level of the post-synaptic proteins were unchanged [18]. Here, we aimed to expand that study and further explore the distribution of presynaptic proteins across the entire human hippocampus and to investigate the possible causes of the proteomic impairment in OML. We selected five presynaptic proteins, i.e. CPLX1, STX1A, SYT1, SYNGR1 and VAMP2, which play important roles in neurotransmission. We performed immunofluorescent staining to assess their expression level assessed across 10 different sub-regions of the hippocampus, categorized into molecular and neuronal layers.

### Heterogeneous distribution of key presynaptic proteins in healthy controls

All selected antibodies showed a strong IR in the neuropil but not in the cell soma such as the densely packed dentate granule cell layer, confirming the enrichment of these proteins in synapses. Interestingly, the five proteins displayed a region-specific distribution which allowed a clear visualization of the OML/IML border: CPLX1, STX1A and SYT1 displayed a stronger IR in the OML whereas SYNGR1- and VAMP2-IR appeared to be more abundant in the IML (Fig. 1). Ramos-Miguel et al. have also detected higher levels of CPLX1 in the OML than in the IML [37]. Albeit both IML and OML contains mainly excitatory inputs to GC, the afferent input is, to a large extent distinct and arises from mossy cells of the CA4 (called associational/commissural projections) and EC layer II, respectively. Thus, it is plausible that the distinctive spatial distribution of the presynaptic proteins occurs as a result of different synapses exhibiting different molecular compositions [34].

### OML-specific presynaptic failure within the hippocampus

The labeling intensity of CPLX1, STX1A, SYT1 and SYNGR1 was significantly reduced in the OML of AD compared to control cases (Fig. 1), corroborating our previous proteomics findings [18].. Interestingly, these reductions were specific to the OML region since all nine other molecular and neuronal sub-regions we examined were preserved, at least in the AD cases examined herein (Fig. 2). Previous studies based on immunohistochemistry reported a decrease in synapsin and synaptophysin in OML, however, detailed analyses of the distribution of these presynaptic proteins between hippocampal sub-regions were missing [21, 25]. In agreement with our study, a decreased ratio of OML/IML levels of synaptic proteins has previously been reported in AD brain, although the potential changes in staining intensity changes in the IML itself was not clearly stated [30, 39]. This specificity is intriguing when considering that our cohort consists of AD cases with late stages of AD pathology (Braak V-VI and amyloid C; Table 2) and that perforant path afferent input from the EC does not solely project to the OML but also to CA fields. Thus, more than a global failure of the perforant path, our results highlight the specific vulnerability of the pathway projecting to the OML in AD, with specific biological mechanisms for this selective vulnerability not yet revealed.

### Possible homeostatic compensation in AD hippocampus

While no profound alterations were detected in the levels of CPLX1, SYT1, SYNGR1 and VAMP2 in any of the other studied hippocampal sub-regions of AD patients, the plasma membrane protein STX1A showed increased immunostaining in the CA4 (Fig. 2). A possible explanation for this finding could be that STX1A levels undergo a compensatory increase in these regions (possibly via increased collateral branching or sprouting) in order to compensate for the reduced input that dentate granule cells receive from the EC layer II, which is evident by the reduced STX1A level in the OML. As opposed to our findings, in a few proteomics study, decreased levels of STX1A were reported in AD brains [7, 32, 58]. However, these studies have used homogenates prepared from bulk tissue of AD brains, and would not be able to tease out cell- and layer-specific discrete changes as the ones observed in sub-hippocampal regions in the current study. The increase in STXA1 staining in the AD cases is clearly visible (Fig. 2C) but only extends through one layer, and therefore would be hard to discern using whole tissue homogenates.

### Specific alteration of OML neurotransmitter release proteins in AD

We next investigated whether the post-synaptic compartment was also altered as expected in the hypothesis of full synapses collapse. When the components of the postsynaptic compartment were investigated, we did not find any alterations in the levels of the postsynaptic density protein SHANK2 and the dendritic marker MAP2 in AD (Fig. 3), which were in line with our previous proteomic findings [18]. Furthermore, the density of dentate granule cells, using NeuN staining, showed no oss of NeuN staining in AD compared to controls. Taken together, our findings suggest that postsynaptic compartments in the OML are rather intact and the deficit is specifically presynaptic.

However, quantification of post-synaptic densities using structural methods such as electron microscopy indicated that they decrease in AD condition [40, 42, 44]. Differences between the amount of synaptic protein amounts (as assessed in the present study) and the number of structurally defined synaptic compartments (as assessed by EM) could account for the divergence in results. Indeed, structural analyses have reported that synapse loss is associated with an increase in the length of synaptic appositions (indicating larger synapses) [44], possibly as a compensation by a synaptic homeostasis mechanism. Therefore, the total amount of synaptic proteins may not change in most synaptic fields. This is however not the case of the OML of the DG, where the decrease in the IR of presynaptic proteins, together with the reduction assessed by our previous proteomic approach [18], advocates for a marked reduction specific to presynaptic proteins. This gives support to the notion of a preferential presynaptic failure in OML of AD brain [3].

Among the proteins we investigated, CPLX1 was the one with the lowest AD/Control ratio both in the present study and in our previous proteomic study [18] performed on different human cohorts, emphasizing the importance of this protein in AD pathogenesis. In agreement with this, previous reports showed that higher levels of specific presynaptic proteins, including CPLX1, were associated with better cognitive function [20, 57]. Before, or in addition to the neurodegeneration occurring in AD, specific alterations in the metabolism of proteins involved in presynaptic plasticity may well contribute to the cognitive impairment.

### Neurodegeneration versus retrograde degeneration

The question of the order of the different pathological events occurring during AD progression is important and has not yet been fully answered. For example, to know whether the presynaptic failure occurs before or as a consequence of neurodegeneration is crucial because it will impact the design of future therapeutical approaches. Compelling neuropathological evidence has shown a dramatic and early loss of EC neurons in AD, particularly in layer II, which correlates with cognitive impairment in AD [16, 24]. Hence, one theory proposes that the degeneration of EC neurons leads to the loss of perforant path afferents and therefore to synaptic loss and reduced synaptic protein-IR in the terminal zone of the perforant path. Supporting this notion, it has been demonstrated that following an EC lesion synaptophysin-IR was reduced in the OML in rat brain [8]. Unfortunately, we were not able to investigate whether the reduction in presynaptic proteins correlated with a loss of EC neurons in layer II since the brain sections in our cohorts were too posterior with neurons in the para-hippocampal cortex not distributed in islands as expected for EC layer II.

We could nevertheless test if the presynaptic changes at the OML were related to axonal degeneration whether this depends or not on somatic degeneration. We assessed possible structural alteration of the axonal projections in OML by investigating the levels of the MBP (for myelinated axons), SMI-312 (for phosphorylated neurofilaments M and H) and the commonly used axonal marker Tau. Despite the advanced stage of the AD cases, we did not find significant alterations in the levels of these three markers in the OML (Fig. 4). This observation is apparently inconsistent with a massive degeneration of EC neurons from layer, a scenario in which the number of projecting afferents should be reduced proportionally to the neuronal loss. One possible explanation is that the afferent input from EC only represent a minority of axons passing through the OML or that axonal projections coming from non-degenerated areas compensate for the loss of axons within the perforant path.

An alternative hypothesis is that axonal projections from EC layer II are not massively degenerated. This possibility offers a straightforward explanation for the unchanged labelling intensities of the axonal markers between AD and controls. It also clarifies why all presynaptic proteins are not altered at the same extent between AD and controls [18]. Indeed, if the reduction in presynaptic proteins was directly resulting from a loss of afferent inputs to the OML, similar AD/Control ratio between the presynaptic proteins detected would have been expected. As previously discussed by Terry et al [52, 53], the loss of neocortical synapses correlates with cognitive decline much better than cell counts and this synaptic loss occurs earlier than death of the neuronal soma. This observation is the basis for the hypothesis of retrograde degeneration which states that synaptic dysfunction precedes axonal degeneration and neuronal death. This hypothesis is supported by a recent observation that changes in synaptic proteins precede neurodegeneration markers in prodromal AD [27]. The data that we collected here are consistent with a retrograde degeneration in which presynaptic proteins are reduced but the axons are not yet degenerated and which, within the hippocampus, would specifically start in the OML.

A retrograde degenerative mechanism could also explain the intriguing observation that the genetic deletion, or the loss of function (using neurotoxins), of several presynaptic proteins leads to neurodegeneration. For example, loss of Munc18 that prevents neurotransmitter release leads to a rapid neurodegeneration [13, 19, 55]. In a similar way, genetic deletion of CSPα revealed that it cooperates with α-synuclein and SNAP25 to prevent neurodegeneration [10, 14, 46] and use of botulinum neurotoxins revealed a direct role of STX1A and SNAP-25 in neuron survival [5, 35]. Thus, there is a possibility that massive reduction in presynaptic proteins, as seen in the perforant path projecting to the OML, leads to the loss of the projecting neurons in the entorhinal cortex. The resolution of the cause-consequence relationship between presynaptic failure and somatic degeneration will be one of the next challenges for research on AD, using studies in populations who are vulnerable for AD.

### Homogeneous burden in amyloid plaques and dystrophic neurites across the hippocampus

We searched for a possible pathological correlate of the presynaptic alteration in OML by investigating the extent of the AD-hallmarks in this region compared to the other hippocampal sub-regions. We labelled amyloid plaques with an antibody against Aβ1-16 and observed neuritic and diffuse plaques in all hippocampal regions in AD whereas amyloid plaques were absent in controls (Fig. 5). Notably, the area covered by amyloid plaques was not larger in the OML compared to other molecular layers of the hippocampus (Fig. 5b). In OML, the presynaptic alterations were not proportional to the amyloid burden (Fig. 5c).

In an effort to specially search for a cause of an axonal or presynaptic impairment we focused our attention on the presence of neurites immuno-reactive for phosphorylated-Tau. Indeed, a role of pathological form of tau in the impairment of presynaptic proteins has been recently exemplified by the observation that pathological Tau binds to synaptic vesicles [26, 59] leading to a smaller pool of active vesicles and to decreased synaptic transmission [59]. We observed that neurites expressing phosphorylated-Tau were remarkably more abundant in AD compared to controls (Fig. 5d). Interstingly, neurites expressing phosphorylated-Tau were arranged sometimes in islands, for example in the IML (Fig. 5d) likely corresponding to dystrophic neurites around senile plaques [1]. The area covered by neurites expressing phosphorylated-Tau was not larger in OML compared to other molecular layers of the hippocampus (Fig. 5d). Moreover, in OML, the presynaptic alterations were not proportional to the burden in phosphorylated-Tau (Fig. 5e).

In summary, our investigation distinctly highlights the projections to the OML, i.e. the perforant path from EC layer II to dentate granule cells, as a specific vulnerable pathway in AD. Our findings also emphasize that postsynaptic compartments in the OML are rather intact, giving support to the notion of specific presynaptic failure in AD. These findings might lead to novel and more targeted therapy options for the early stages of AD.

## Abbreviation list

AD: Alzheimer’s disease
CA: Cornu ammonis
CPLX1: Complexin-1
CPLX2: Complexin-2
EC: Entorhinal cortex
IML: Inner molecular layer
LM: Stratum lacunosum-moleculare
IR: Immuno-reactivity
LUC: Stratum lucidum
MAP2: Microtubule-associated protein 2
MBP: Myelin basic protein
MOPP: Molecular layer perforant path-associated cells
NEUN: Neuronal nuclei
OML: Outer molecular layer
PMI: Postmortem interval
RAD: Stratum radiatum
SHANK2: SH3 and multiple ankyrin repeat domains protein 2
SNARE: Soluble NSF-attachment protein receptor
STX1A: Syntaxin-1A
SYNGR1: Synaptogyrin-1
SYT1: Synaptotagmin-1
VAMP2: Vesicle-associated membrane protein 2

## Acknowledgements

The *postmortem* human brain tissue for this study was supplied by the Netherlands Brain Bank (Amsterdam, the Netherlands). We thank all the donors for the tissue used in this study. Imaging was performed at the Bordeaux Imaging Center, a service unit of the CNRS-INSERM and Bordeaux University, member of the national infrastructure France BioImaging (ANR-10-INBS-04).

## Author contributions

H.H., T.J.S., S.F. and G.B. designed the research. T.J.S. performed immunofluorescence experiments. H.H. did densitometry measurements and data analysis. H.H., S.F., T.J.S and G.B. wrote the manuscript. C.M., A.-C.G., L.T.O. and B.W. contributed to the writing of the manuscript. The manuscript was read, edited and approved by all authors.

## Materials and methods

### Post-mortem human brain tissues

Formalin-fixed paraffin-embedded hippocampal sections (5 μm thick) from eight sporadic AD and seven non-demented control cases were obtained from the Netherlands Brain Bank. AD cases were clinically diagnosed according to previously published research criteria [12] and pathologically confirmed by the presence of amyloid plaques and neurofibrillary tangles. On the other hand, control cases showed no sign of dementia and presented no or little pathological alterations including a few plaques and tangles. The demographic characteristics of AD and control cases are shown in Table 2. All donors or their next-of-kin gave informed consent. The establishment of the Netherlands Brain Bank was approved the Independent Review Board of VU University Medical Center in Amsterdam, the Netherlands (2009/148).

### Immunofluorescence

Hippocampal sections were first deparaffinized in xylene (≥ 98.5 %, AnalaR NORMAPUR® ACS, Reag. Ph. Eur. analytical reagent) and then rehydrated using the following wash steps: 99 %, 95 % and 70 % ethanol, and distilled water. Subsequently, heat-induced antigen retrieval was done using citrate buffer (10 mM citric acid (Sigma Aldrich), with 0.05 % Tween-20 (Sigma Aldrich), pH 6) at 110 °C for 30 min inside a pressure cooker (Biocare Medical). Slides were then washed in PBS with 0.05% Tween-20 (PBS-T) and were subsequently blocked with normal goat serum (Thermo Fisher Scientific) for 20 min at room temperature (RT). Slides were incubated with the primary antibodies listed in Table 3 for 2 hours at RT. In order to reduce autofluorescence, slides were then incubated with 1x TrueBlack® Lipofuscin Autofluorescence Quencher (Biotium), which was diluted in 70% ethanol, for 5 min at RT. Subsequently, sections were washed with 1x PBS for three times, which was followed by the incubation with the secondary antibodies (anti-rabbit, anti-mouse and anti-guinea pig IgG conjugated to Alexa 488 or Alexa 568, Thermo Fisher Scientific, 1:500 dilution) for 2 hours at RT. Sections were washed with 1x PBS for three times and incubated with either 5 µM of Methoxy X04 (Tocris Bioscience) for 20 min at RT or 300 nM of DAPI for 10 minutes at RT. Finally, sections were washed in 1x PBS three times and in distilled water, cover-slipped using Fluoromount™ (Southern Biotech) and air-dried overnight at RT. In order to avoid inter-experimental variability, sections from all AD and control cases were stained with a certain antibody at the same time.

**Table 3:** The list of primary antibodies used in this study.

| Antibodies | Source | Dilution |
| --- | --- | --- |
| CPLX1 antibody, rabbit polyclonal | #10246-2-AP, Protein Tech | 1:500 |
| STX1A antibody, rabbit polyclonal | #ANR-002, Alomone | 1:1000 |
| SYNGR1 antibody, mouse monoclonal | #103011, Synaptic Systems | 1:500 |
| SYT1 antibody, mouse monoclonal | #105011, Synaptic Systems | 1:500 |
| VAMP2 antibody, mouse monoclonal | #104211, Synaptic Systems | 1:500 |
| MBP antibody, rabbit monoclonal | #ab216668, Abcam | 1:500 |
| Aβ 1-16 antibody (DE2), mouse monoclonal | #MAB5206, Sigma Aldrich | 1:500 |
| Phospho-Tau antibody, mouse monoclonal | #MN1060 Thermo fisher scientific | 1:500 |
| SMI-312 antibody, Purified anti-neurofilament marker, mouse polyclonal | #837904, BioLegend | 1:500 |
| SHANK2 antibody, guinea pig polyclonal | #162204, Synaptic Systems | 1:500 |
| Tau-1 antibody, mouse monoclonal | #MAB3420, Sigma-Aldrich | 1:500 |
| MAP2 antibody, mouse monoclonal | #M4403, Sigma-Aldrich | 1:500 |
| NeuN antibody, rabbit polyclonal | #ABN78, Merck | 1:500 |

Additionally, we assessed the specificity of the fluorescent signal coming from the secondary antibodies by performing several control experiments. In the first experiment, we tested the presence of non-specific fluorescence signal that can be caused by the secondary antibody itself, and incubated consecutive hippocampal sections from a healthy control case with either primary (i.e. STX1A and SYT1) and secondary antibodies (i.e. Alexa 488 and Alexa 568) or only secondary antibodies. In the second experiment, we tested the efficiency of TrueBlack to remove the endogenous autofluorescence coming from the tissue (Jordá-Siquer et al. unpublished results).

### Imaging and image analysis

Slides were loaded into the semi-automated Nanozoomer 2.0HT slide scanner (Hamamatsu) and scanned using FITC (green), TRITC (red) and DAPI channels at 20x magnification and fixed additional lens 1.75X. Image acquisitions and settings were kept constant for all sections stained with a certain antibody. Additionally, to avoid biases in fluorescence intensity, all AD and control cases were also scanned together. Gamma correction of 1 was applied to all images in order to reach linearity in fluorescence signal and images were then exported as tiff files using the NDP.view2 software (Hamamatsu) and analyzed using ImageJ Fiji (NIH).

Semi-quantitative assessment of fluorescence pixel intensity of CPLX1, STX1A, SYT1, SYNGR1 and VAMP2 stainings was carried out in the following sub-regions of the hippocampus: molecular layers of the dentate gyrus (IML and OML), molecular layers of CA3 (CA3-LUC, CA3-RAD, CA3-LM), molecular layers of CA1 (CA1-RAD and CA1-LM) as well as the neuronal layers like CA4, CA3 and CA1. Mean pixel intensities of the area of interest were measured in a blinded fashion. The background, which is the signal intensity measured outside the tissue, was subtracted from each case. In order to explore changes occurring in the presynaptic and postsynaptic compartments, we did densitometric analysis of three axonal markers (MBP, SMI-312 and Tau) as well as of a postsynaptic density marker (SHANK2) and a dendritic marker (MAP2) in the OML and the adjacent IML. Additionally, we measured the neuronal density of the dentate granule cells, which were normalized to the total surface area (mm^2^), by using NeuN and DAPI stainings in a blinded fashion. Similarly, amyloid and phosphorylated Tau burden were determined by measuring the mm^2^ area in each of the synaptic regions of interest normalized to the total surface of the analyzed region.

### Statistical analysis

Demographic characteristics between AD and control groups as well as semi-quantitative densitometric assessments were analyzed in GraphPad PRISM 7.0 (San Diego, CA). Two-tailed Student’s t-test was used to assess whether age, brain pH and PMI significantly differ between control and AD groups, while Mann-Whitney test was used to assess the statistical significance for gender, ApoE status, Braak and amyloid stages between the two groups. Statistical analysis of semi-quantitative densitometric assessment of fluorescence pixel intensity was done using either two-tailed Student’s t-test or Mann-Whitney test, depending on whether the data follows a normal distribution or not, which was checked by Kolmogorov-Smirnov test. The presence of outliers was also checked by the ROUT (default) method. In all statistical comparisons between AD and control groups, p-value < 0.05 was considered statistically significant.

## Figure Legends

**Supplementary Figure 1.**
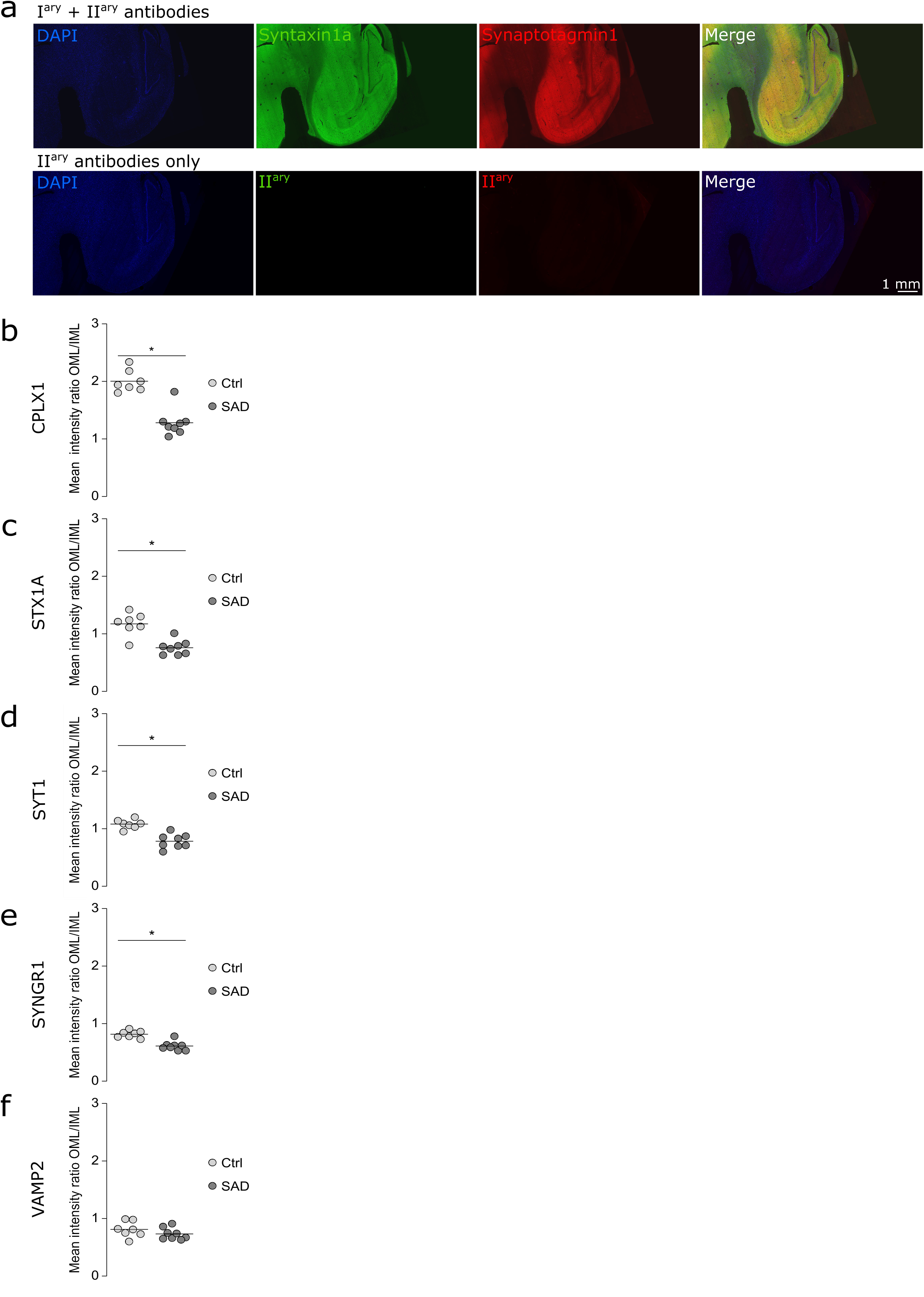
Impaired OML/IML intensity ratio for presynaptic proteins. (a) Controls of fluorescent immuno-labeling in human hippocampal sections. In the upper panels, the sections are stained for DAPI, syntaxin1a and Syt1. In the lower panels, only secondary antibodies were incubated in addition to DAPI. The fluorescent immuno-labelings are specific to the primary antibodies. (b-f) Scatter plots of the mean pixel intensity OML/IML ratio of the fluorescent labellings respectively for CPLX1, STX1A, SYT1, SYNGR1, VAMP2. The ratios were significantly decreased in AD cases by 36 % for CPLX1 (t-test, p < 0.0001), 35 % for STX1A (p = 0.0003), 28 % for SYT1 (p = 0.0001) and 25 % for SYNGR1 (p < 0.0001). The ratio for VAMP2 showed was unaltered.

**Supplementary Figure 2.**
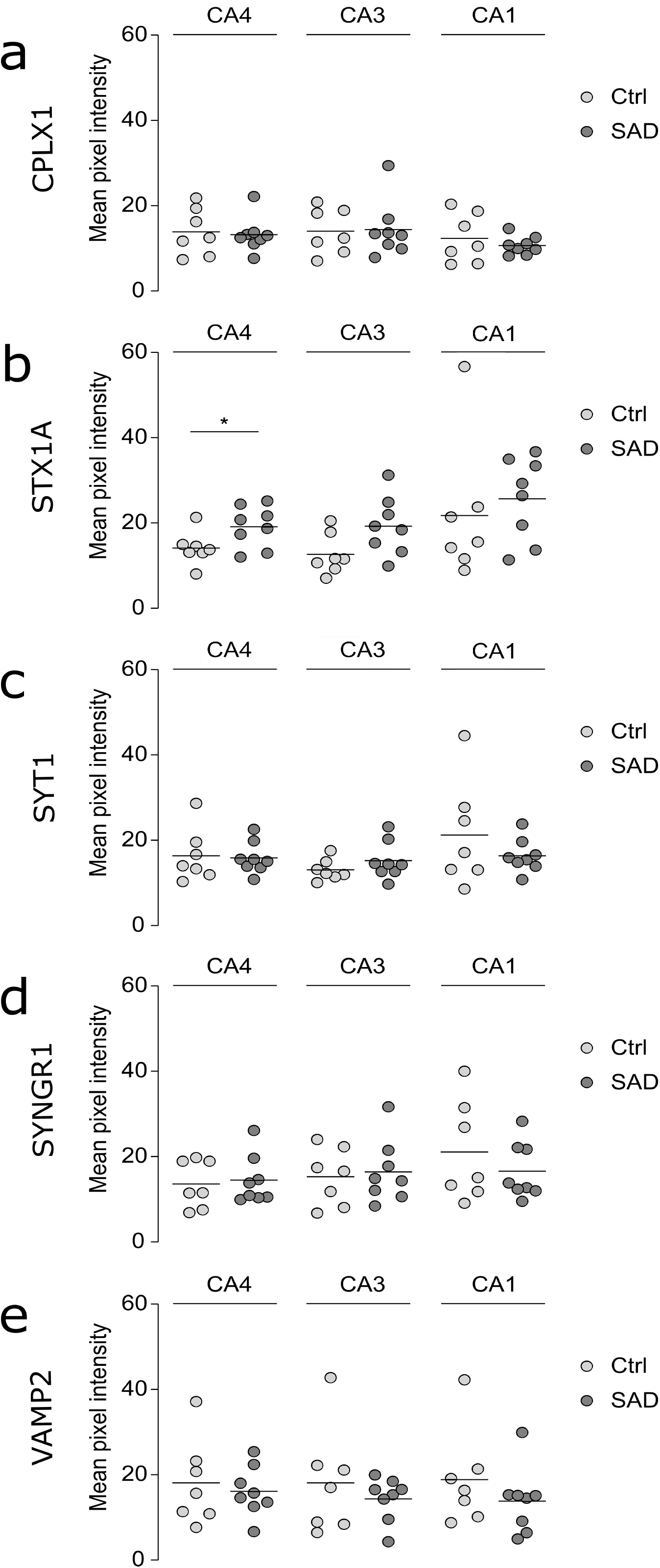
Preserved level of presynaptic proteins in various neuronal layers of the hippocampus. (a-e) Scatter plots of the mean pixel intensity of the fluorescent labellings in CA3, CA1 and CA4 respectively for CPLX1, STX1A, SYT1, SYNGR1, VAMP2. The fluorescence pixel intensities were not altered in these regions except for STX1A which was increase in AD CA4.

